# Reducing maladaptive behavior in neuropsychiatric disorders using network modification

**DOI:** 10.1101/2024.08.02.606383

**Authors:** Nicholas Timme

## Abstract

Neuropsychiatric disorders are caused by many factors and produce a wide range of symptomatic maladaptive behaviors in patients. Despite this great variance in causes and resulting behavior, we believe the maladaptive behaviors that characterize neuropsychiatric disorders are most proximally determined by networks of neurons and that this forms a common conceptual link between these disorders. Operating from this premise, it follows that treating neuropsychiatric disorders to reduce maladaptive behavior can be accomplished by modifying the patient’s network of neurons. In this proof-of-concept computational psychiatry study, we tested this approach in a simple neural network model that produces aversion-resistant alcohol drinking – a key maladaptive behavior associated with alcohol use disorder. We demonstrated that it was possible to predict personalized network modifications that substantially reduced maladaptive behavior without inducing side effects. Furthermore, we found that it was possible to predict effective treatments with limited knowledge of the model and that information about neural activity during certain types of trials was more helpful in predicting treatment than information about model parameters. We hypothesize that this is a general feature of developing effective treatment strategies for networks of neurons. This computational study lays the groundwork for future studies utilizing more biologically realistic network models in conjunction with *in vivo* data.

## Introduction

A central goal of neuroscience is the discovery of new treatments for neuropsychiatric disorders like addiction, depression, schizophrenia, anxiety, and attention-deficit/hyperactivity disorder (ADHD). Patients with neuropsychiatric disorders present with a wide range of symptomatic behavior[1] and many causal factors are thought to contribute to these disorders[2–4]. Despite this heterogeneity, we believe it is helpful to focus our attention at the level of networks of neurons because the maladaptive behavior associated with these neuropsychiatric disorders are most proximally produced by the network of neurons in the patient’s brain. In other words, we believe maladaptive behaviors such as consuming a drug of abuse, inattention to the task at hand, or suffering a hallucination can be viewed as the manifestation of malfunctioning networks of neurons. Adopting this view, the question of how to reduce maladaptive behavior can then be recast as what parameters of the patient’s network of neurons should be modified to reduce maladaptive behavior (Figure 1). In this way, the question of how to treat maladaptive behavior can be reduced to questions about neural populations, such as what populations of neurons are too active or not active enough, what connections are too strong or too weak, and so forth.

**Figure 1:**
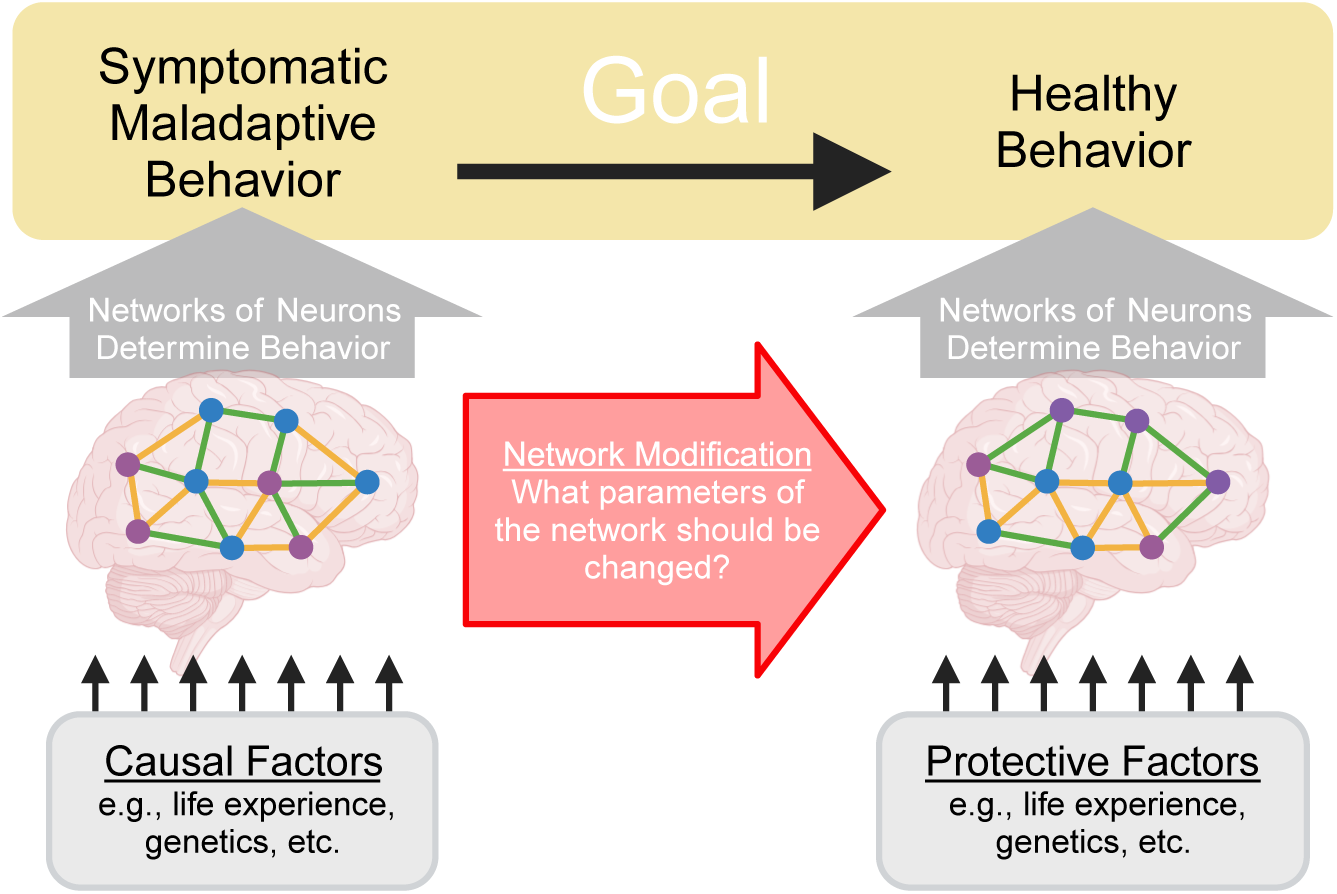
Treating neuropsychiatric disorders can be recast as modifying network parameters. A central goal of neuroscience is to treat neuropsychiatric disorders by reducing symptomatic maladaptive behavior and increasing healthy behavior. Crucially, these behaviors are determined by activity patterns within the patient’s networks of neurons. Therefore, to alter behavior, it is necessary to identify and manipulate network parameters. Causal factors for the disorder can be conceptualized as influencing this system by altering the network of neurons in certain ways to produce malfunctions that then drive symptomatic maladaptive behavior.

We recognize that our network centric viewpoint may seem controversial because it appears to discard known causal factors for neuropsychiatric disorders and existing treatments. On the contrary, we wish to emphasize that we believe it is vital to understand how causal factors – such as social factors, genetic vulnerabilities, past trauma, or other life experiences – drive these disorders. We believe these causal factors can fit within the network centric viewpoint as factors that alter the network of neurons, ultimately leading to the aberrant network behavior that determines maladaptive behavior associated with the disorder (Figure 1). Furthermore, we believe existing treatments – such as pharmacological treatments and cognitive behavioral therapy – can be viewed as processes that modify the network of neurons to reduce maladaptive behavior, even though these treatments are not typically conceptualized in those terms.

We believe explicitly focusing on networks of neurons possesses a key advantage when studying neuropsychiatric disorders. These disorders are driven by multicausal factors that are personalized to each patient[2–4]. However, previous studies have primarily focused on identifying isolated individual causal factors that contribute to neuropsychiatric disorders[5–8]. Understanding how multiple causal factors interact in each individual patient is a daunting task that is not well suited to these traditional reductive investigatory methods. It may be the case that manipulating the brain to reverse one causal factor found in isolation is insufficient to effectively treat the disorder[9, 10] or produces unintended off target effects (i.e., side effects)[11, 12]. Despite the highly varied nature of patient specific multicausal factors, all of these factors converge on the network of neurons. Therefore, by understanding the neural cause of symptomatic maladaptive behavior, it may be possible to substantially improve treatment. In other words, it may be the case that the universe of neural mechanisms that drive symptomatic maladaptive behavior is smaller than the universe of causal factors that can produce those neural mechanisms. Thus, focusing on the network of neurons and how it should be modified to reduce symptomatic maladaptive behavior may be easier than understanding complex interactions between multiple causal factors. While the general view that neuropsychiatric disorders originate from malfunctioning networks of neurons is implicit in a great deal of neuroscientific research, we have not found previous research explicitly formulating neuropsychiatric treatment as network modification. That said, we believe this approach generally falls within the domain of computational psychiatry[13, 14], which seeks to leverage computational approaches to model neural behavior[15, 16] and predict treatment of neuropsychiatric disorders[17]. Some studies have sought to use neural networks to understand interactions between symptoms (so called, “symptom networks”)[18]. Alternatively, connectionist[14, 19, 20] approaches have sought to use neural networks to model the generation of the maladaptive behavior associated with neuropsychiatric disorders[21]. In line with these connectionist approaches, we will utilize neural network models that produce maladaptive and healthy behaviors to understand the effects of manipulating network parameters on behavior and as a test subject for predicting treatment.

The program we propose is also closely related to network control theory, which seeks to develop methods to control network behavior, typically by identifying which parts of the network can control the rest of the network[22]. This work has been pursued in generalized types of networks[23], as well as in models of brain networks[24, 25]. In brain networks, the goal of this research is to produce behavior changes by controlling neural activity in certain key nodes in the network[26], using tools such as transcranial magnetic stimulation[27]. Work has also been done on structural control theory to examine how changing the structure of networks can produce different behavior, though often in abstract terms[28–30]. Overall, our goal of modifying behavior by changing network parameters is well aligned with network control theory and structural control theory.

Taken together, previous work in computational psychiatry and network control theory provides numerous connection points in terms of methods and motivations for our program of neural network modification. However, we are unaware of a previous example explicitly involving the treatment of neuropsychiatric disorders by modifying network parameters. Thus, we believe a feasibility study with this goal that uses a simple computational model would help to build intuition about this method and drive development of more advanced techniques that could be applied *in vivo*. We will focus on a particular example of maladaptive behavior associated with a particular neuropsychiatric disorder: aversion-resistant alcohol drinking in alcohol use disorder. This key symptom of alcohol use disorder involves continuing to drink alcohol despite negative consequences[1]. Using a computational model will provide us with a best-case scenario in terms of knowledge of the parameters that control the network. This will allow us to explore the feasibility of using network modification techniques to reduce maladaptive behavior. At this early stage of development, we chose to use a simple model that is not biologically realistic and is not bound by patient data, other than with regards to exhibiting general symptomatic behavior associated with the disorder. This model is most similar to other connectionist models that do not seek to directly relate network models to actual brain networks. Furthermore, our focus at this stage is not to explicitly connect network modification with existing or potential clinical treatments, though this will be an important consideration in later stages of developing this method.

Our primary goal is to determine the feasibility of using network modification techniques to reduce maladaptive behavior. We hypothesize that (1) it will be possible to identify key parameters in the model that can reduce maladaptive behavior without producing side effects, (2) that it will be possible to predict effective treatments that are individualized to each specific model, and that (3) it will be possible to create these predictions with limited knowledge of the model. We will begin by building a simple neural network model that produces aversion-resistant alcohol drinking behavior. Next, we will use traditional systems neuroscience techniques to characterize this model. Finally, we will treat the model to reduce maladaptive behavior and examine techniques to make personalized network modifications. This study will lay the groundwork for future studies utilizing more biologically realistic models and *in vivo* data.

## Results

### Task and Network Structure

We created a simple neural network model that produces aversion-resistant alcohol drinking (Figure 2 A) to easily explore the role played by network parameters in aversion-resistant drinking and how these parameters can be changed to reduce maladaptive behavior. Prior to selecting the model structure, we designed a task with four possible inputs and four possible action outputs (Figure 2 B) that allows for specificity to aversion-resistant drinking. We intentionally limited the stimuli and behavioral outputs to a few simple options to ease interpretation of results. The model was trained and tested on trials, during each of which the model was presented with one of six possible combinations of inputs (i.e., trial types) and the model selected one of the four possible output actions. The target input to action mappings for the six trial types used in training (Figure 2 C) were chosen because they demonstrate proper danger processing, appropriate eating behavior, and aversion-resistant or aversion-sensitive alcohol drinking. For instance, an immediate danger cue (e.g., a large oncoming truck) should produce an output/action of escape. A food availability cue (e.g., a cheeseburger) should produce an output/action of eat, unless it appears that the food is not safe due the presence of a consumption risk cue (e.g., a nearby bottle of poison). An alcohol availability cue (e.g., a glass of wine) should produce an output/action of drink. However, if a consumption risk cue (e.g., needing to drive) is present along with the alcohol availability cue, aversion-sensitive models should choose to wait and aversion-resistant models should choose to drink. We chose to construct models that exhibit maladaptive behavior selectively to allow for the assessment of side effects produced by network parameter manipulation. In this case, side effects were incorrect responses on non-aversion-resistance related trials, such as immediate danger avoidance trials.

**Figure 2:**
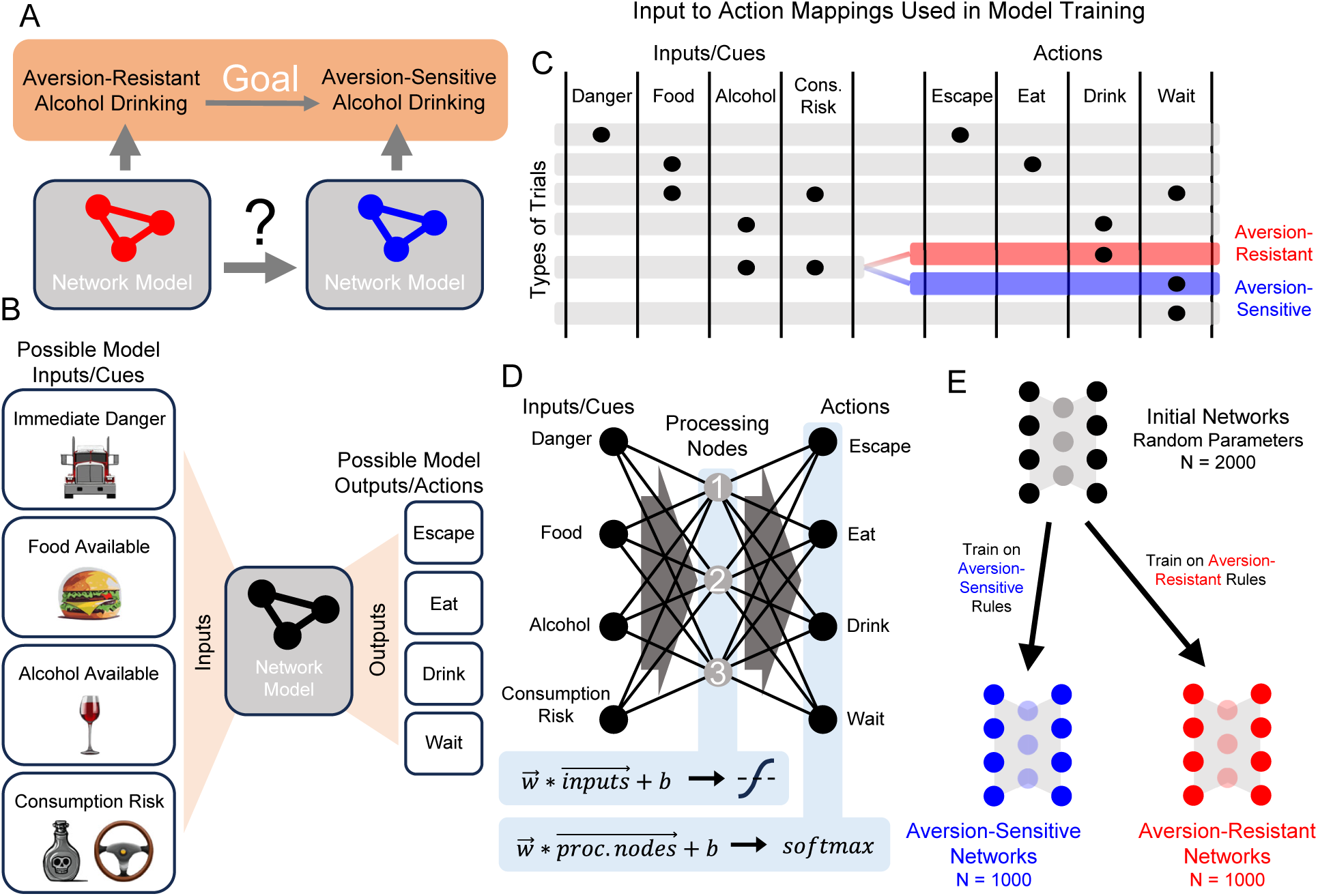
Demonstration of neuropsychiatric disorder treatment via network modification in a simple computational model of aversion-resistant alcohol drinking. **(A)** Consider two computational models that exhibit aversion-resistant and aversion-sensitive alcohol drinking. Our goal is to modify the aversion-resistant model so that it becomes aversion-sensitive. **(B)** Four possible inputs/cues (immediate danger, food available, alcohol available, and consumption risk) were supplied to the model and the model selected one of four possible output actions (escape, eat, drink, or wait). **(C)** Six patterns of inputs (i.e., trial types) were used, each of which possessed a target output that was used in model training. Aversion-resistant and aversion-sensitive models were trained on identical input/output mappings, except for trials where both alcohol availability and consumption risk inputs were active. In these trials, aversion-resistant models were trained to select drink and aversion-sensitive models were trained to select wait. **(D)** A simple three-layer feedforward neural network was capable of learning both types of rules. This network possessed one hidden layer with 3 processing nodes and non-linear activation functions. **(E)** 2000 initially random networks were trained on either aversion-sensitive or aversion-resistant rules to produce 1000 networks of each type.

We chose to use a small feedforward neural network to study the behavior associated with these aversion-resistant or aversion-sensitive behavior rules (Figure 2 D). We chose this model because it was the smallest feedforward neural network with a hidden layer that could learn the aversion-resistant and aversion-sensitive rules. (See Discussion Section for more information about the realism of the model.) We refer to the hidden layer nodes as “processing” nodes because they do not receive stimuli or determine behavioral outputs. Initially random networks were trained on either the aversion-resistant or aversion-sensitive rules to greater than 95% accuracy using 1000 randomly selected input/output training combinations (Figure 2 E). This model possessed 31 parameters including 12 connection weights from input nodes to processing nodes, 12 connection weights from processing nodes to output nodes, 3 biases associated with processing nodes, and 4 biases associated with output nodes. Note that connections weights could be positive (excitatory) or negative (inhibitory).

### Characterizing the Network with Systems Neuroscience Techniques

Following network training, we used systems neuroscience techniques to examine the role played by the processing neurons in the model networks (Figure 3 A-E). First, we silenced each processing node and examined the resulting changes in network behavior (Figure 3 A, B). This method is similar to ablation/lesion experiments[31] or studies of behavioral changes following localized brain injuries. Silencing processing node 1 resulted in deficits in correct escaping behavior, silencing processing node 2 resulted in deficits in correct eating behavior, and silencing processing node 3 resulted in deficits in drinking behavior (Figure 3 B). In addition, silencing processing node 3 in aversion-sensitive networks also reduced correct eating behavior.

**Figure 3:**
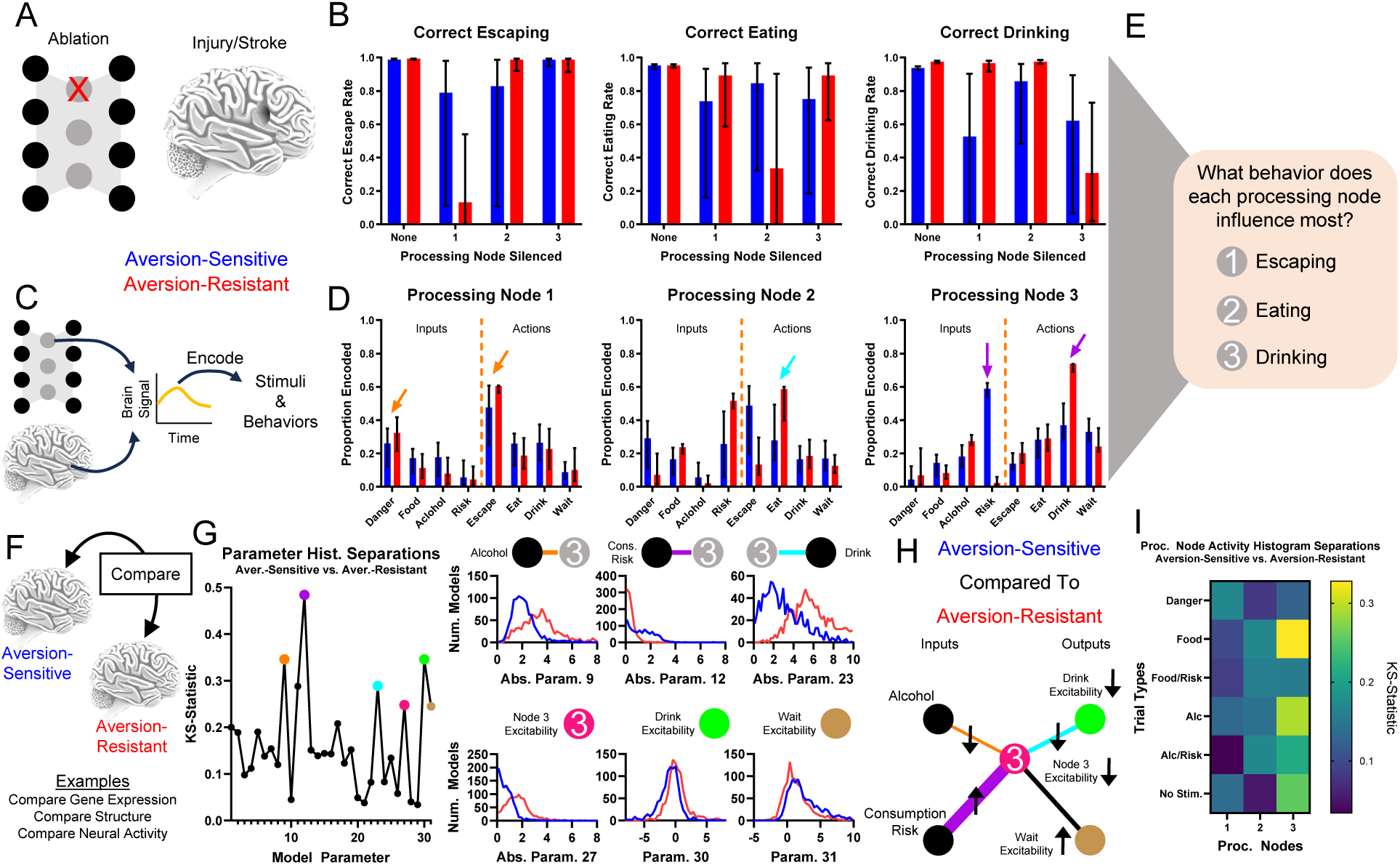
Traditional systems neuroscientific techniques elucidate processing node roles and identify potential treatment targets. Examining behavioral changes following damage to a population of neurons (e.g., ablation, stroke, injury) **(A)** and studying encoding of stimuli and behaviors by brain signals (e.g., electrophysiology, fMRI) **(C)** are common methods to determine the role of neural populations in behavior. **(B)** Silencing (artificially setting node activity to 0) resulted in key deficits in correct escaping (node 1), correct eating (node 2), and correct drinking behavior (node 3) for the three processing nodes, respectively. **(D)** Neural encoding by each node of the stimuli and behavioral outputs indicated that node 1 best encoded signals related to escaping immediate danger (orange arrows), node 2 best encoded signals related to eating (cyan arrow), and node 3 best encoded signals related to drinking (purple arrows). Though, note that node 3 encodes almost no information about danger and much more information about drinking in aversion-resistant networks. **(E)** Taken together, (A-D) indicate that node 1 was most involved with escaping, node 2 was most involved with eating, and node 3 was most involved with drinking. **(F)** Brain features (e.g., gene expression, brain structure, neural activity) of patients and healthy controls are often compared to identify the cause of maladaptive behavior. **(G)** The distributions of parameter values for aversion-sensitive and aversion-resistant models were compared. Six of the parameters with the largest differences in distributions were related to processing node 3 (distributions of parameter absolute values shown in some cases to collapse across excitatory and inhibitory connections). **(H)** Specifically, aversion-sensitive models possessed elevated drive from the consumption risk input, elevated excitability of the wait output, weakened drive from the alcohol input and to the drink output, and weakened excitability of processing node 3 and the drink output (portion of the model with only these parameters shown). **(I)** The distributions of processing node activity values for each type of trial for aversion-sensitive and aversion-resistant models were compared. Activity values differed most in processing node 3. (Median and interquartile range shown in all bar graphs, N = 1000 for aversion-resistant and aversion-sensitive models.)

We also examined encoding by processing nodes of input stimuli and behaviors (Figure C, D), which is similar to neural encoding experiments that seek to relate BOLD or electrophysiological signals to stimuli or behaviors. Processing node 1 most encoded danger and escaping, processing node 2 most encoded risk and eating, and processing node 3 most encoded drinking (Figure 3 D). We also found that in aversion-resistant networks, processing node 3 encoded very little information about risk and a great deal of information about drinking. Overall, these analyses indicate that, in general, processing node 1 most influences escaping, processing node 2 most influences eating, and processing node 3 most influences drinking (Figure 3 E).

To further investigate the function of the network, we compared the parameter values and processing node activity levels between aversion-resistant and aversion-sensitive networks (Figure 3 F). This type of comparison is similar to studies that examine differences between patients and healthy subjects in terms of brain structure[32, 33], gene expression[34, 35], or neural activity[36, 37]. We compared the distribution of parameters using the KS-statistic, which assesses how similar the distributions of parameter values were in these two types of models (Figure 3 G). We highlighted six parameters with the large differences that were related to processing node 3. Aversion-sensitive networks had higher connection strengths from the consumption risk input to processing node 3, lower connection strengths from the alcohol input to processing node 3, lower connection strengths from processing node 3 to the drink output, higher excitability (bias) in the wait output, and lower excitability (bias) in processing node 3 and the drink output (Figure 3 H). Thus, aversion-sensitive networks functioned such that the alcohol stimulus stimulated processing node 3 less, which was less excitable, which drove the drink output less, which was also less excitable. We also compared the distributions of processing node mean activity values between aversion-resistant and aversion-sensitive networks for each of the six different types of trials (Figure 3 I). In alignment with differences found in parameters, we found differences in activity were highest in processing node 3.

### Network Treatment

To move beyond simply characterizing the networks and their differences, we examined ways to treat aversion-resistant networks by modifying their parameters. First, we retrained the aversion-resistant networks using the aversion-sensitive rules (Figure 4 A). As expected, this treatment dramatically reduced aversion-resistant drinking (trials where the model selected drink when the alcohol and consumption risk inputs were active) without increasing side effects (errors on all types of trials) (Figure 4 B). While this result is encouraging, it required retraining the whole network and adjusting all parameters. To assess whether it was necessary to retrain the whole network, we instead sequentially imposed only the largest parameter changes from retraining (Figure 4 C). We found that network performance improved with additional parameter changes, but it was necessary to impose several parameter changes to reach aversion-sensitive network performance levels.

**Figure 4:**
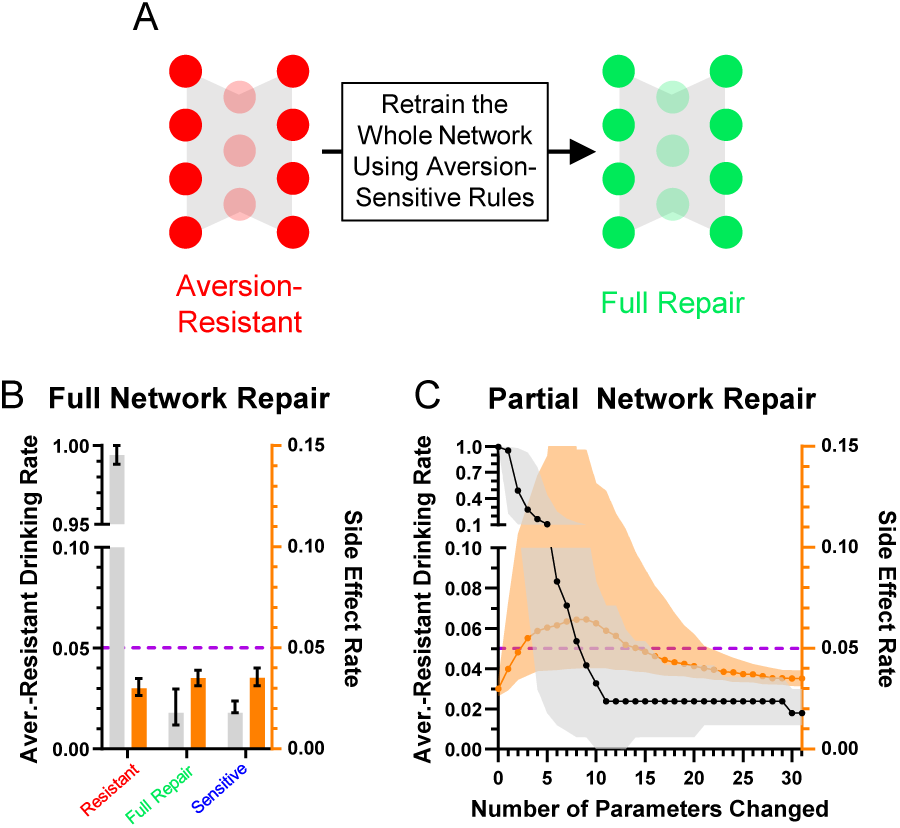
Aversion-resistant models can be cured by retraining the network. **(A)** Aversion-resistant models were first treated by retraining the whole model with the aversion-sensitive rules. **(B)** As expected, this process dramatically reduced aversion-resistant drinking without inducing side effects. **(C)** Imposing parameter modifications found during whole network retraining in decreasing order of change in magnitude demonstrated that the largest magnitude parameter changes reduced aversion-resistant drinking, but many parameter changes were necessary to fully reduce aversion-resistant drinking levels to those found in aversion-sensitive models. (Median and interquartile range, N = 1000 for aversion-resistant and aversion-sensitive models.)

While this method demonstrated that it was not necessary to change every network parameter to effectively treat aversion-resistant drinking, we wondered if another method would yield better improvement with fewer parameter changes. To investigate this possibility, we exhaustively searched through each parameter to minimize the aversion-resistant drinking and side effects (Figure 5 A). The best parameter change found using this search method produced substantially reduced aversion-resistant drinking in comparison to the largest magnitude change from the whole network retraining or the average parameter modification for the best search parameter (Figure 5 B), demonstrating the power of personalized treatments even in this simple model. We then examined how performance improved for each of the networks’ 31 parameters (Figure 5 C). In particular, parameter 12 (CoR3: connection weight from consumption risk input to processing node 3) often reduced aversion-resistant drinking dramatically without increasing side effects and was the parameter that, when modified, most frequently reduced total error (aversion-resistant drinking rate plus side-effects rate) the most (Figure 5 D). Note that this parameter was one of the parameters with the largest difference between aversion-resistant and aversion-sensitive models (Figure 3 G), but that other parameters identified in that analysis were not as capable of reducing aversion-resistant drinking behavior alone. We examined the relationship between the original CoR3 parameter value in the aversion-resistant network and the new value found via search to reduce aversion-resistant drinking (Figure 5 E). We found a weak relationship between the two parameter values, indicating that new CoR3 parameter values could not be easily estimated based solely on the CoR3 parameter value in aversion-resistant networks. We also compared model performance following CoR3 modification to the average model performance following modification of the seven parameters with the largest difference between aversion-resistant and aversion-sensitive network (Figure 5 F). We found that the search method reduced aversion-resistant drinking substantially more compared to the parameters identified with the traditional approach.

**Figure 5:**
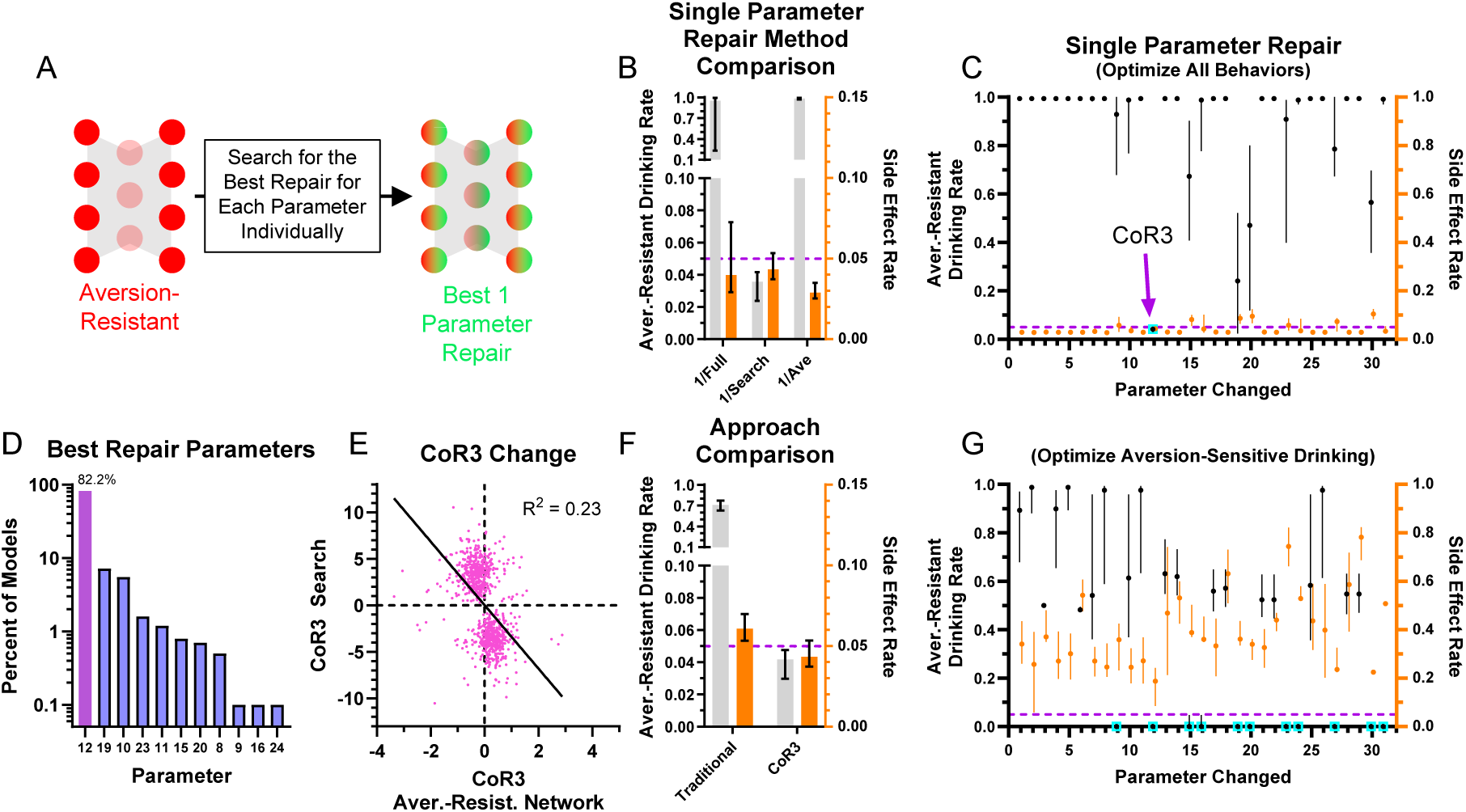
Most aversion-resistant models could be effectively treated by modifying one parameter. **(A)** Rather than retraining the whole network, each individual parameter was modified to search for the best possible reduction in aversion-resistant drinking for each parameter. **(B)** Imposing the single best parameter change from the search process (1/Search) reduced aversion-resistant drinking substantially more than imposing the largest magnitude parameter change from whole network retraining (1/Full), without substantially increasing side effects. For comparison, imposing the average parameter modification for the parameter found via search did not reduce aversion-resistant drinking (1/Ave) (i.e., the non-personalized approach). **(C)** Aversion-resistant drinking and side effect rates found for each parameter modification individually using the exhaustive search. Despite identifying several parameters with substantial differences between aversion-resistant and aversion-sensitive model (Figure 3 G), only modification of parameter 12 (connection weight from the consumption risk input to processing node 3 (CoR3), cyan box) could reduce aversion-resistant drinking to low levels for nearly all models while maintaining low side effect rates. **(D)** Histogram of parameter changes that were found to produce the best treatment in terms of reduction in aversion-resistant drinking rate and side-effect rate (CoR3: 82.2% of networks). **(E)** The CoR3 parameter value found via search was weakly related to the original CoR3 parameter value in the aversion-resistant network. **(F)** Aversion-resistant drinking was substantially lower in models following CoR3 modification than the average aversion-resistant drinking following modification of the seven parameters that were found to be most different between aversion-resistant and aversion-sensitive models (Figure 3 G). **(G)** The lowest aversion-resistant drinking rates (regardless of side effect rates) for each parameter individually using the exhaustive search. Note that aversion-resistant drinking could be substantially reduced by modifying numerous parameters (cyan boxes) if side effects were ignored. (Median and interquartile range, N = 1000 for aversion-resistant and aversion-sensitive models.)

We also examined how effective parameter changes were if only considering aversion-resistant drinking and ignoring side effects (Figure 5 G). We found that it was possible to substantially reduce aversion-resistant drinking by modifying numerous parameters, but these changes resulted in increased side effects.

### Network Treatment Prediction

We have shown that it is possible to find individual parameters such that their modification can substantially reduce aversion-resistant drinking without increasing side effects. However, finding these parameter changes was time consuming and required computationally expensive searches across all network parameters. In the future, with more complex and biologically realistic network models, this strategy may become prohibitive. So, we examined whether it was possible to predict parameter changes from the network parameters or processing node activity patterns using partial least squares (PLS) methods (Figure 6, see Methods Section for a detailed description of the PLS models).

**Figure 6:**
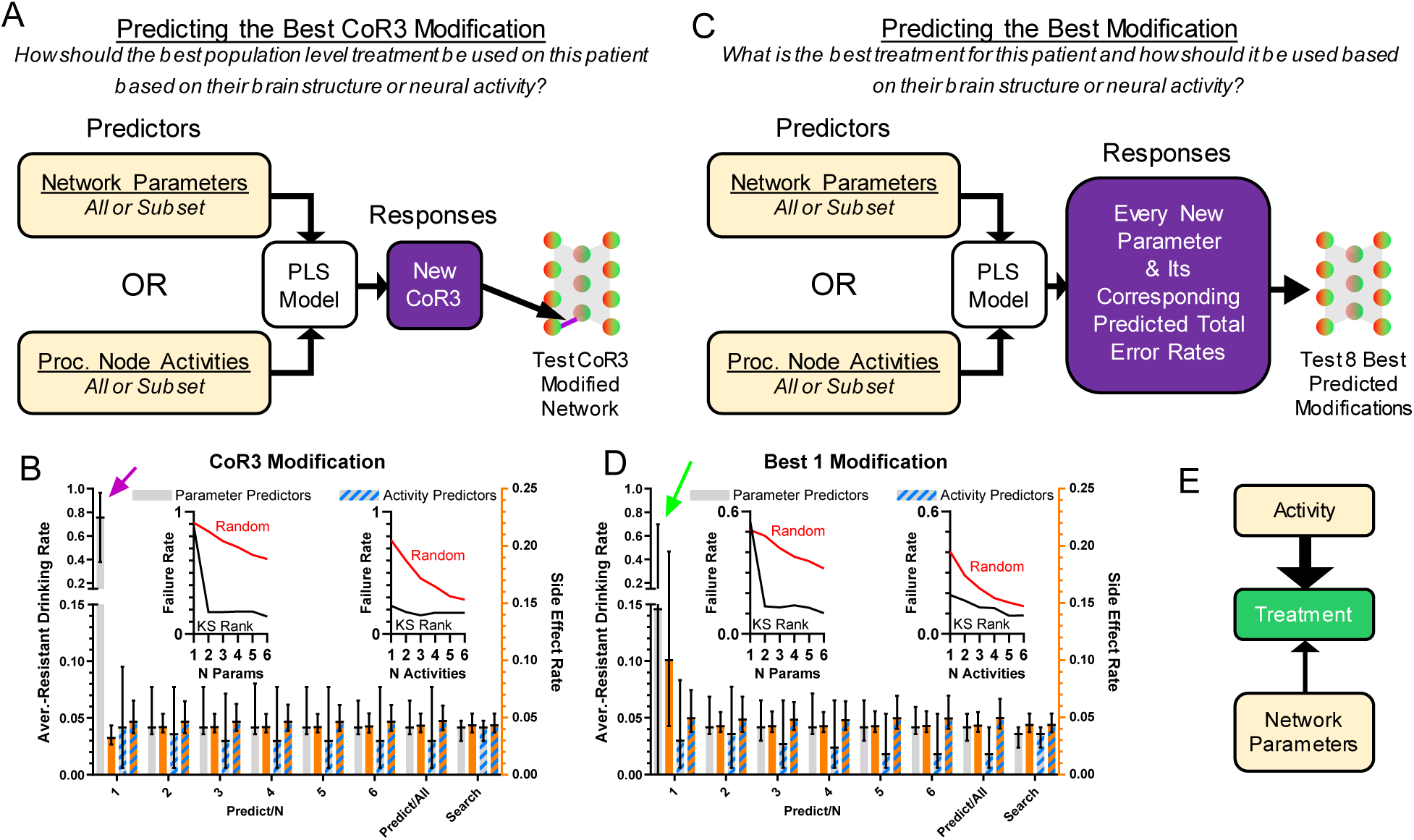
Effective treatments could be predicted from model structure and neural behavior. **(A)** Treatment by modification of CoR3 was predicted using a PLS model based on network parameters or processing node activity. Three types of predictors were used: all parameters, the average processing node activities on all trial types, or subsets of 1-6 parameters/activities (in descending KS-statistic rank (Figure 3 G and I)). CoR3 values found via search (Figure 5 C) were used as responses for PLS fitting. New CoR3 values produced by the PLS model were imposed on each model and tested. **(B)** Model behavior with a PLS predicted CoR3 update using parameters or processing node activity as predictors. Activity predicted successful treatment better than network parameters when prediction was performed with 1 activity value or parameter (magenta arrow). **(C)** Instead of predicting CoR3 values, new PLS models were created to predict the best new value for each parameter and the corresponding total error rate (aversion-resistant drinking rate plus side effect rate) for each parameter change. The eight parameter changes with the lowest predicted error rate were then tested and the parameter change with the lowest error rate was selected. **(D)** Model performance with a PLS model predicted parameter update using parameters or processing node activity as predictors. Activity predicted successful treatment better than network parameters when prediction was performed with 1 activity value or parameter (green arrow). For all types of PLS models using subsets of highly separated parameters (Figure 3 G) or activity (Figure 3 I) values as predictors, the treatment failure rates (proportion of models with aversion-resistant drinking rates over 0.1) decreased rapidly with the inclusion of more predictors and were generally lower than models with randomly chosen parameters/activities as predictors (B and D insets). **(E)** In general, treatment was better predicted by network activity than network structure, particularly when limited data about the model was available. (Median and interquartile range, N = 1000 models.)

Given the success of CoR3 modification in the parameter search, we first constructed PLS models to predict new values of CoR3 that would reduce aversion-resistant drinking without increasing side effects (Figure 6 A). In these PLS models, model predictors were either network parameters or the average processing node activities on the six different types of input trials. Furthermore, we tested whether it would be possible to make effective CoR3 predictions with only subsets of the parameters or activities. To do so, we ranked the parameters and the activities by their separation (KS-statistic) between aversion-resistant and aversion-sensitive models (Figure 3 G and I) and we tested 1-6 of the parameters or activities in descending order of separation. The parameter and activity with the largest KS-statistic were, respectively, Cor3 and the average activity of processing node 3 on safe eating trials. The human treatment analog for this treatment prediction would be predicting how to use the best population level treatment in a patient based on their brain structure or neural activity.

We found that PLS models could predict CoR3 modifications that successfully treated the aversion-resistant models to a degree similar to the search method (Figure 6 B). Furthermore, we found that prediction treatment performance improved substantially with the inclusion of more parameters as predictors and that this performance reached asymptote near the search method performance at 5 parameters. The value of CoR3 was the parameter with the highest separation (i.e., Predict/1 in Figure 6 B), so note that knowledge of CoR3 alone was not sufficient to accurately predict the best CoR3 treatment (see Figure 5 E). We also found that inclusion of parameters with high aversion-resistant vs. aversion-sensitive separations reduced treatment failure rates (proportion of networks with aversion-resistant drinking rates over 0.1) faster than randomly selected parameters. We found that treatment performance was substantially better using activity as the predictor relative to network structure when only one predictor variable was used.

Though treatment by CoR3 modification was found to be successful in many models, we utilized a second type of PLS model to instead predict the best modification (Figure 6 C). This PLS model used the same types of predictors as the previous model. However, instead of predicting just the new Cor3 value, all new parameter values and their corresponding total error rates (aversion-resistant drinking rate plus side effects rate) were predicted. After predicting the new parameter changes and their corresponding error rates, the eight parameters with the lowest predicted error rates were tested in the aversion-resistant model and the one with the lowest error rate was selected. The human treatment analog for this type of treatment prediction would be predicting what would be the best treatment in a patient and how to use it based on their brain structure or neural activity.

Similar to the PLS model used to predict the best CoR3 change, predicting the best parameter to change produced low aversion resistant drinking rates that were generally improved by adding more predictors (Figure 6 D), especially when using parameters as predictors. Furthermore, we found that activity values were again better able to predict treatment in comparison to network structure, especially when including only one predictor variable. Overall, using both PLS treatment prediction methods, we found that network activity predicted treatment better than network structure when few variables were used for prediction (Figure 6 E).

Lastly, we examined networks for which the search of all single parameter changes did not produce a network with sufficiently low aversion-resistant drinking behavior (Figure 7 A). About 4.2% of networks did not achieve aversion-resistant drinking rates of less than 10%, the threshold we set for insufficient aversion-resistant drinking reduction (Figure 7 B). In these networks, we employed an exhaustive search of all pairs of parameters and found that it was possible to reduce aversion-resistant drinking yet further in these networks (Figure 7 C).

**Figure 7:**
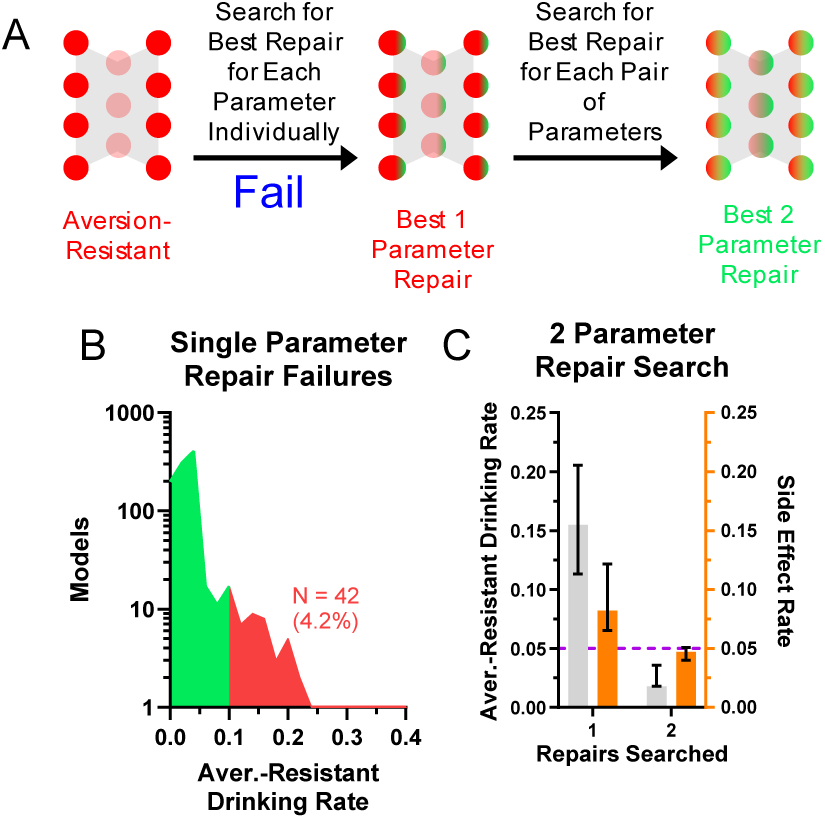
Networks that were not effectively treated with one modification could be treated with two modifications. **(A)** For a small subset of networks, an exhaustive search of individual parameter changes did not reduce aversion-resistant drinking sufficiently. **(B)** Approximately 4.2% of networks did not achieve aversion-resistant drinking rates below 10% with only one parameter modification. **(C)** An exhaustive search of all two parameter changes in these networks did substantially reduce aversion-resistant drinking. (Median and interquartile range shown in all bar graphs, N = 1000 models.)

## Discussion

We demonstrated that it was possible to effectively treat aversion-resistant drinking in a computational model by modifying network parameters, that these treatments were personalized to each individual model, and that these treatments could be predicted using computationally efficient predictions with limited information about the model. This provides proof of concept evidence that reducing maladaptive behaviors associated with neuropsychiatric disorders by modifying networks of neurons is a viable treatment for neuropsychiatric disorders.

### Treatment Prediction with Limited Data

We found that processing node activity was better able to predict effective treatments when only limited information is known about the model (Figure 6 B and D). We hypothesize that this is a general feature of the network modification treatment approach for situations with limited data. This result is intuitive because node activity is what ultimately determines behavior, so knowledge of node activity will likely be more helpful in determining how to change parameters to correct aberrant node activity than information about one of the numerous parameters that goes into determining node activity. This hypothesis, if true, has important implications for testing decisions and the development of new tests because it indicates that methods that measure neural activity (e.g., electrophysiology and fMRI) will be best positioned to predict treatment *in vivo*.

### Limitations and Advantages of the Model

In this study we examined a specific type of neuropsychiatric disorder and a simple model of one of its associated maladaptive behaviors. This model is not an adequate model of the human brain, but we wish to emphasize that this was not the goal for this project. Rather, we sought to create a model that balanced detail with generalizability in a way where treatment predictions in a neural network that governed behavior could be tested and interpreted. The model we created strikes this balance appropriately because it is defined by numerous parameters that can be related to networks of neurons (e.g., connection weights, biases), but it could be generalized to many different types of maladaptive behaviors, it is conceptually tractable, and it could incorporate analyses of side effects. We make no claim that this model is superior to other existing or potential models of aversion-resistant drinking and so we performed no comparison with other models. Rather, the goal of our model was to serve as a test bed for the network modification program.

While the model we created met the goals of this project, it possesses many shortcomings, especially regarding behaviors and phenomena specific to alcohol use. For instance, our model did not incorporate information about the temporal structure of drinking (both within and between drinking episodes), the pharmacology of alcohol, the effects of alcohol history on brain function, or the role of specific vs. generalized deficits in decision-making. Furthermore, it did not incorporate various other psychological factors involved in alcohol use disorder, such as craving, stress, or negative affect. Also, the stimuli presented to the model and actions performed by the model were dramatically simplified in comparison to the rich array of stimuli and behaviors associated with alcohol use disorder in humans. Finally, the simplicity of the model rendered it impossible to connect model behavior with actual *in vivo* data from pre-clinical or clinical studies, other than with regards to the fact that the model produced behavior that generally aligns with the aversion-resistant drinking phenotype that is central to the alcohol use disorder diagnosis[1]. We believe all of these are important factors in developing a more thorough model of alcohol use disorder, and we aim to include them in the future. Other researchers have included some of these features in models of alcohol use disorder[38–40], but these other models were not structured as neural networks making it difficult to understand how modifications to the models could translate to potential treatments. In this study, by ignoring these complicating factors, we were able to improve generalizability and simplicity, which we felt was paramount to our research program at this stage.

### Future Research

In the future, we will improve the techniques developed herein (Figure 8). Next, we will aim to improve the biological realism of the neural network model to include features such as time, learning, realistic neural connectivity and firing properties, and realistic neurotransmitter dynamics, as well as other neuropsychiatric disorders. A crucial question in these studies will be how to predict effective treatments in more complex models that will likely possess many more parameters.

**Figure 8:**
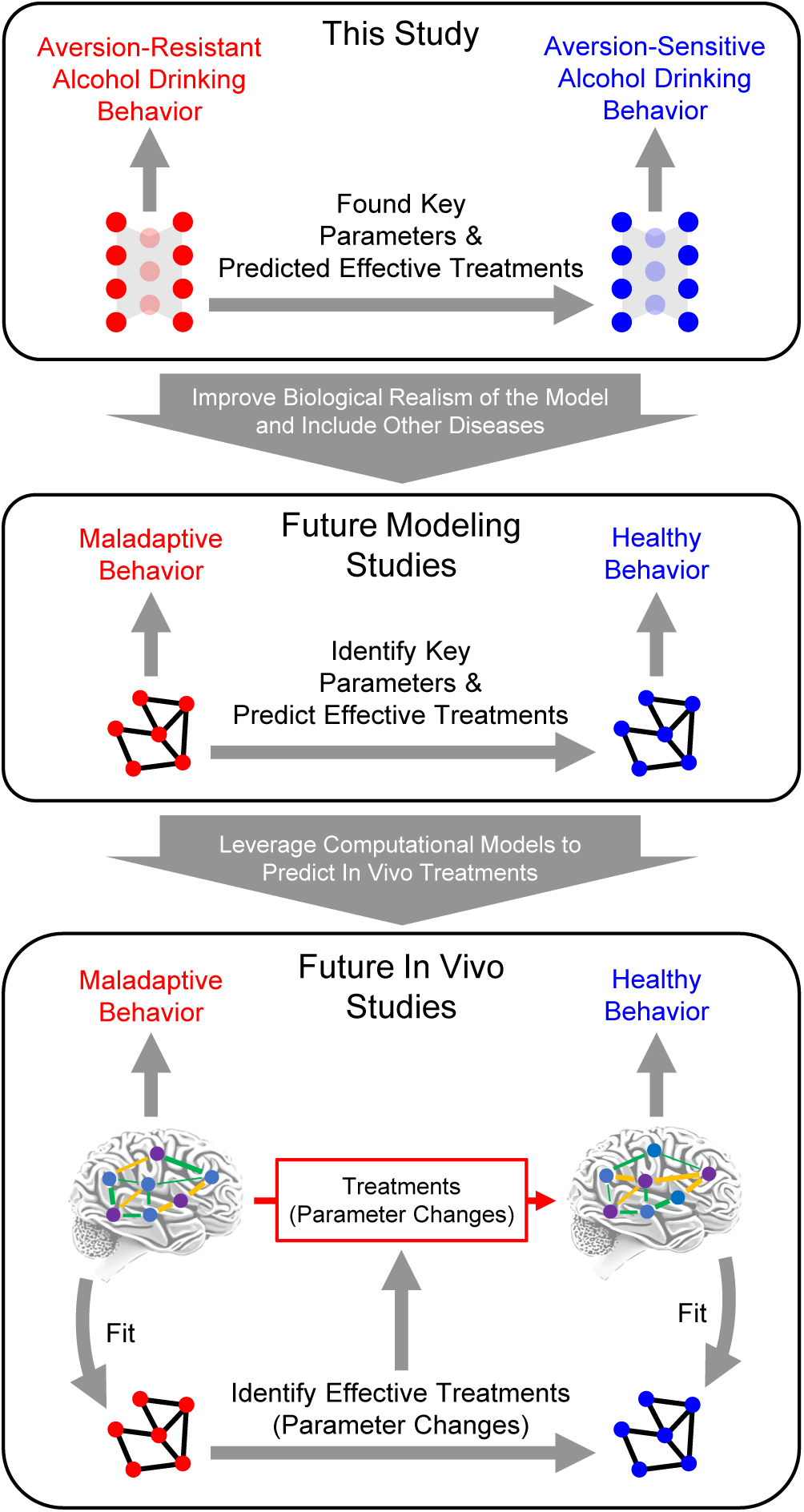
Future computational psychiatry research plan. This study utilized a simple feedforward neural network as a model system to demonstrate the feasibility of reducing maladaptive behavior by modifying neural network parameters. Key parameters were found in this model and effective treatments were predicted based on model structure and model behavior. In the future, it will be vital to expand this research to include other neuropsychiatric disorders and more biologically realistic models. Following that work, it will be possible use computational psychiatry approaches and leverage computational models to predict *in vivo* treatments by fitting computational models to *in vivo* networks, using those models to test treatments in silico, and then predicting *in vivo* treatments.

Following these improvements in silico, we will apply these methods *in vivo* by fitting a computational model to the *in vivo* data, then testing modification methods in the computational model, and then testing whether those manipulations resulted in effective treatments when applied *in vivo*[13, 16]. By leveraging neural network models in concert with *in vivo* data, it will be possible to generate and test falsifiable hypotheses about how *in vivo* neural networks function[20]. These studies will likely be first conducted in pre-clinical rodent models, but our hope is that these techniques will be usable in human subjects. Importantly, these methods may indicate that a certain type of network parameter modification is necessary, but there may be no feasible technique to do so in humans. If so, this result will drive development of new methods to manipulate neural populations in humans. Furthermore, these methods can be applied to existing treatments to better understand how they operate, if they can be improved, and how they could be personalized to individual patients.

### Characterizing Neural Function

We utilized several traditional system neuroscience techniques to characterize the role played by processing nodes and the network parameters in the model. The processing node silencing manipulations and neural encoding analyses provided a picture of distinct roles for the three processing nodes. However, processing node silencing often produced minor deficits in several behaviors and most processing nodes encoded some information about all inputs and outputs. Thus, though certain processing nodes appeared to most contribute to certain behaviors, the roles played by these nodes were complex and not precisely delineated.

Examining parameter differences between aversion-resistant and aversion-sensitive networks highlighted several parameters as potential targets for treatment. These differences produced a logical explanation for how aversion-sensitive networks produced more wait outputs rather than drink outputs during aversion-resistant drinking test trials. However, when these parameters were individually manipulated, only one was found to substantially reduce aversion-resistant drinking without inducing side effects. Numerous other parameters could be modulated to reduce aversion-resistant drinking, but at the cost of increased side effects. We suggest this demonstrates a significant concern regarding neural manipulation experiments in pre-clinical rodent models in particular. Frequently, these studies use optogenetics or chemogenetics to increase or decrease activity in a certain neural population and then assess behavioral changes that result. While these studies typically utilize controls, such as locomotor behavior to ensure that decreases in a maladaptive behavior are not simply due to a general decrease in activity, it is extremely difficult to assess all possible side effects of the neural activity manipulation *in vivo*. As a result, this type of study may produce a false confidence that the neural population that was manipulated can be best understood as controlling the maladaptive behavior, when really the manipulation of that population may just be capable of disrupting the maladaptive behavior and many other untested behaviors. Modeling approaches such as those used in this study are more capable of quickly testing all possible side effects and identifying targeted manipulations that minimize side effects.

Overall, we are concerned that these types of systems neuroscience analyses lean heavily on the assumption that individual parts of the brain or any network should have clear roles that we can understand[41, 42]. Rather, there is substantial evidence that understanding synergistic neural behavior across multiple neural populations is necessary to understand complex behaviors[43–45]. Furthermore, it has been shown that traditional systems neuroscience tools like those used here are not capable of providing thorough explanations for complex information processing systems and that non-linear dynamical models, such as neural network models we propose to use in the future to capture *in vivo* network behavior, are better suited to this goal[46]. Even though we showed that modification of one parameter in particular (CoR3) was an effective treatment, it was not possible to determine how to modify this parameter using only information about that parameter. In other words, even in the case of this simple network, it was necessary to incorporate information from multiple parts of the network to determine the correct parameter modification. In more complicated networks, we hypothesize that the need to utilize information from multiple parts of the network will become even more critical and that it will be necessary to modify more than one parameter to effectively treat neuropsychiatric disorders. Furthermore, when more complex models are used, it may be necessary to employ advanced techniques like artificial intelligence and machine learning[47, 48] to predict effective treatments instead of the PLS methods used herein.

## Acknowledgements

We would like to thank Christopher Lapish, Will Barnett, Cherish Ardinger, Kari Haines, Maribel Hernandez, Meredith Bauer, David Swygart, and Christina Gremel for their helpful comments throughout the development of this project.

This work was supported by the National Institutes of Health (www.nih.gov) (NMT: AA028265). The funder played no role in study design, data collection and analysis, the decision to publish, or preparation of the manuscript.

## Methods

### Data and Analysis Code Availability

All Matlab code necessary to reproduce the data and analysis discussed in this article, along with the Prism file used to generate figures, is available online[49].

### Model Networks

Each model network possessed four input nodes, three hidden nodes in a single layer, and four output nodes. In the manuscript we refer to the hidden layer nodes as “processing” nodes. Strictly feedforward connections existed from the input layer to the hidden layer and from the hidden layer to the output layer. The connection weight from input layer node *j* to hidden layer node *i* was given by *w*_*i*,*j*,1_ and the connection weight from hidden layer node *j* to output layer node *i* was given by *w*_*i*,*j*,2_.

The activity of input layer node *i* was given by *x*_*i*,1_ and was set for each run of the network by a unique combination of four input variables that corresponded to the four possible stimuli (danger, food, drug, and consumption risk). Each stimulus could be off (0) or on (1). Noise, in the form random gaussian variables with a standard deviation of 0.2 was added to these input values.

The activity of hidden layer node *i* was given by Eq. (1), where *b*_*i*,1_ was the bias for hidden layer node *i* and the tansig activation function was used.

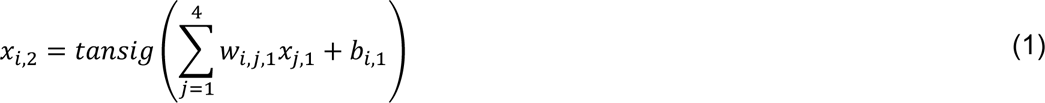

The activity of output layer node *i* was given by Eq. (2), where *b*_*i*,2_ was the bias for output layer node *i* and the softmax activation function was used.

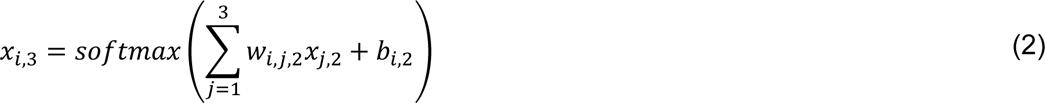

Due to the softmax activation function, the output layer node activities were bound between 0 and 1 for all inputs. The output behavior of the network was set according to the output node with the highest value.

The networks were trained using one of nine randomly selected training algorithms[50] and a training data set of 1000 randomly generated input/output combinations set according to the rules for aversion-resistant and aversion-sensitive behavior (Figure 2). Random networks were created and trained on either aversion-resistant or aversion-sensitive training data. Aversion-resistant networks were stored and then retrained on the aversion-sensitive data to produce fully retrained networks. If a network produced error rates of less than 5% for the given training rules, it was accepted and used in further analyses. Random network creation and training continued until 1000 aversion-resistant and 1000 aversion-sensitive networks were produced.

Following training, a testing data set of 1000 randomly generated input combinations was run through all networks. Output node activities and hidden layer node activities were recorded for these test input combinations. All remaining analyses utilized these activities as well as the parameters (*w*_*i*,*j*,1_, *w*_*i*,*j*,2_, *b*_*i*,1_, and *b*_*i*,2_) associated with each network.

### Information Theory Analysis

The encoding of network inputs and outputs by hidden layer activities was calculated to characterize the behavior of the hidden layer neurons[51]. For inputs, the encoding was calculated as the difference between the total joint mutual information with all inputs and the joint mutual information with the other inputs in order to remove interactions between inputs. Furthermore, the information was normalized by dividing by the entropy of the hidden layer neuron. The input and output states were already discretized to be 0 or 1. The hidden layer activities were discretized using four equal counts bins. The encoding of input *j* by hidden layer node *i* was given by Eq. (3), where the mutual information was given by Eq. (4), the entropy was given by Eq. (5), *X*_*i*_ was the set of possible hidden layer activity values (four discretized values), *Y*_[1,4],1_ was the set of all possible combinations of input values, and *Y*_[1,4]−*j*,1_ was the set of all possible combinations of inputs values excluding input *j*.

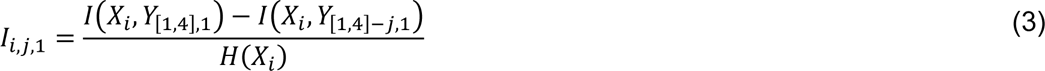

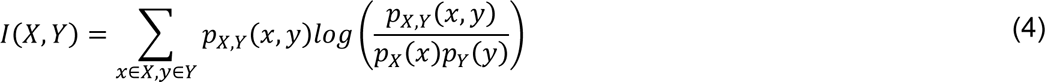

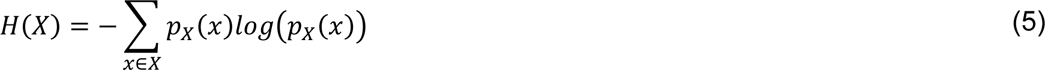

The encoding of output *j* by hidden layer node *i* was given by Eq. (6), where *Y*_*j*,2_ was the set of possible output values of output *j*.

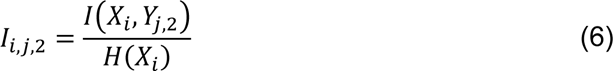

Note that because the outputs were mutually exclusive due to the softmax activation function, it was not possible to remove interactions between outputs.

### Hidden Layer Node Number

Due to the random initial structure of the networks, it was necessary to renumber hidden layer nodes according to their functional role because nodes serving the same role could have been assigned to different numbers in the hidden layer by chance. To do so, we used an unsupervised method to cluster nodes into three functional groups. First, the eight encoding values for each hidden layer node (encoding of four inputs and encoding of four outputs) were subjected to a principal component analysis across all hidden layer nodes from all networks of a given type (aversion-resistant or aversion-sensitive). The first two principal component scores then underwent k-means clusters to detect 3 clusters. The hidden layer nodes for each network were then assigned to one of these clusters based on minimum overall distances in principal component space. The mean cluster encodings were then assigned a label such that the cluster with the highest encoding of the escape output was labeled node 1, then the highest encoding of the eat output was labeled node 2, and the remaining node was labeled node 3. These new numbers were then used for the hidden layer nodes throughout the analysis and the connection weight and bias parameters were adjusted to function with this renumbering.

### Hidden Node Silencing

To investigate the role of hidden layer nodes, each node was systematically silenced or ablated by artificially setting its output value to 0 regardless of its inputs and the test data set was run through the model. The performance was then assessed by calculating the rate of correct task performance on a variety of task scenarios. For instance, for the immediate danger/escape task scenario, the rate of correct escaping was the number of trials where the model selected escape (i.e., the output node with the largest activity was escape) divided by the number of trials where it should have selected escape (i.e., the danger input was active).

### Model Comparison

To investigate the differences between aversion-resistant and aversion-sensitive models, the KS-statistic was calculated for the distributions of parameter values and average processing node activities between aversion-resistant and aversion-sensitive models. Larger KS-statistics indicated that the distributions of parameter values were more different.

### Model Retraining

Following initial training for aversion-resistant rules, models were retrained with aversion-sensitive rules. The performance of the models was assessed by calculating the aversion-resistant drinking rate and the side effect rate. The aversion-resistant drinking rate was the proportion of trials where the alcohol and consumption risk inputs were active and the model selected drink. The side effect rate was the proportion of all trials where the model did not select the correct action (e.g., the model did not select escape when the input was danger).

In addition to assessing the performance of the aversion-resistant, fully retrained network, and aversion-sensitive network, performance was also assessed in models with partial retraining. In these networks, the parameter changes were ordered by the magnitude of the parameter change during retraining. Parameter changes were imposed in order from largest to smallest and the performance of the network was assessed with each additional parameter change.

### Parameter Search

Each parameter in the aversion-resistant models was optimized to reduce overall error on aversion-sensitive rules (i.e., reducing both aversion-resistant drinking and side effects) and just minimizing aversion-resistant drinking. This minimization was accomplished using an iterative random search process of parameter values. On each iteration, 100 random numbers were selected near the current parameter values using a gaussian distribution (initial standard deviation equal to 10, centered on current parameter value). The network was run for each of these parameter values and the value with the lowest error (either on aversion-sensitive rules or on just aversion-resistant drinking) was selected. In the next iteration, the standard deviation of the new random numbers was set as the minimum of 0.5 and the difference between the previous iteration parameter value and the new iteration. This process was continued until the new iteration error improved by less than 1% of the previous iteration error.

For networks that failed to produce a single parameter change that resulted in less than a 10% aversion-resistant drinking rate when considering all aversion-sensitive rules, a second search was performed for each pair of parameters using the same methods as those described for a single parameter search, with one exception. For searches of two parameters, instead of 100 randomly selected new parameter values, there were 100 randomly selected new parameter value pairs for the two parameters being searched.

### Treatment/Parameter Prediction

A partial least squares (PLS) method was used to predict parameter changes using a variety of predictor and response combinations. In all cases, responses for PLS fitting were taken from parameter search results and fitting was performed using data from all models other than the individual model for which PLS was used to predict parameter modification. This implies that the results we obtained for PLS predicted treatments could be applied successfully to new aversion-resistant models without requiring computationally expensive parameter searches in new aversion-resistant models.

Let *A*_*i*,*j*,*k*_ represent the average activity for processing node *j* on trial type *k* (recall, there are six types of trials (see Figure 2 C)) for aversion-resistant model *i*. Let {*A*_*i*,*j*,*k*_} represent the set of *A*_*i*,*j*,*k*_ values for all processing node and trial type combinations. Let {*A*_*i*,*j*,*k*_} represent the set of *A*_*i*,*j*,*k*_ values for the processing node and trial type combinations that had the one through *l* largest KS-statistics between aversion-resistant and aversion-sensitive model (see Figure 3 I). Let *B*_*i*,*j*_ represent parameter *j* for aversion-resistant model *i*. Let {*B*_*i*,*j*_} represent the set of *B*_*i*,*j*_ values for all parameters. Let {*B*_*i*,*j*_} represent the set of *B*_*i*,*j*_ values for the parameters that had the one through *l* largest KS-statistics between aversion-resistant and aversion-sensitive model (see Figure 3 G). Let 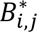 represent parameter *j* for aversion-resistant model *i* found via search that most reduced aversion-resistant drinking and side effects. Let {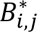} represent the set of 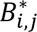 for all parameters. Let 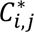 represent the total error rate (aversion-resistant drinking rate plus side effect rate) found in model *i* after imposing parameter 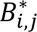. Let {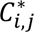} represent the set of 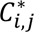 for all parameters.

Two types of PLS models were produced. First, some PLS models attempted to predict the new parameter 12 (CoR3) value for a given network that was found via the search algorithm above. Parameter 12 was the connection weight from the consumption risk input node to hidden layer node 3. It was selected because it was found to reduce aversion-resistant drinking most substantially while simultaneously not increasing side effects. In these PLS models, the response for the PLS model fit for aversion-resistant model *m*: *m* ≠ *i* was {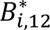} and the combinations of predictors used in different PLS models were subsets of activities 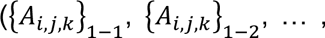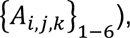, all activities ({*A*_*i*,*j*,*k*_}), subsets of parameters 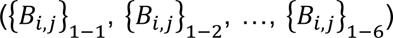, or all parameters ({*B*_*i*,*j*_}). In these PLS fits, each model *i* was a single observation. After fitting each PLS model for model *m* based on one of the sets of predictors, the PLS model was used to predict a new value for parameter 12 based on the corresponding predictors from model *m*. This new parameter was then imposed on the model and its performance was assessed.

Second, some PLS models attempted to predict all parameters found by search and their associated total error rate. In these PLS models, the responses for the PLS model fit for aversion-resistant model *m*: *m* ≠ *i* were {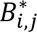} and {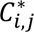} and the combinations of predictors used in different PLS models were subsets of activities 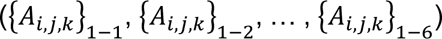, all activities ({*A*_*i*,*j*,*k*_}), subsets of parameters 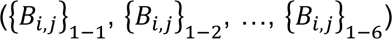, or all parameters ({*B*_*i*,*j*_}). In these PLS fits, each model *i* was a single observation. After fitting each PLS model for model *m* based on one of the sets of predictors, the PLS model was used to predict new parameter and total error rates based on the corresponding predictors from model *m*. Of these new parameters, the eight with the lowest predicted total error rate were each imposed on the model and its performance was assessed. The parameter change that produced the lowest total error rate was then selected and used as the predicted treatment.

## Notes

https://figshare.com/articles/dataset/Data_and_Analysis_Code_Reducing_maladaptive_behavior_in_neuropsychiatric_disorders_using_network_modification/26417524?file=48084769

### Competing Interest Statement

The authors have declared no competing interest.

